# G_n_T motifs: short nucleotide tracts of ≥8bp that can increase T:A→G:C mutation rates >1000-fold in bacteria

**DOI:** 10.1101/2024.12.30.630749

**Authors:** James S. Horton, Joshua L. Cherry, Gretel Waugh, Tiffany B. Taylor

## Abstract

Nucleotides across a genome do not mutate at equal frequencies. Instead, specific nucleotide positions can exhibit much higher mutation rates than the genomic average due to their immediate nucleotide neighbours. These ‘mutational hotspots’ can play a prominent role in adaptive evolution, yet we lack knowledge of which short nucleotide tracts drive hotspots. In this work, we employ a combinatorial approach of experimental evolution with *Pseudomonas fluorescens* and bioinformatic analysis of various *Salmonella* species to characterise a short nucleotide motif (≥8bp) that drives T:A→G:C mutation rates >1000-fold higher than the average T→G rate in bacteria. First, we experimentally show that homopolymeric tracts (≥3) of G with a 3’ T frequently mutate so that the 3’ T is replaced with a G, resulting in an extension of the guanine tract, i.e., GGGGT → GGGGG. We then demonstrate that the potency of this T:A→G:C hotspot is dependent on the nucleotides immediately flanking the G_n_T motif. We find that the dinucleotide pair immediately 5’ to a G_4_ tract and the nucleotide immediately 3’ to the T strongly affect the T:A→G:C mutation rate, which ranges from ∼5-fold higher than the typical rate to >1000-fold higher depending on the flanking elements. Therefore the T:A→G:C hotspot motif is a product of several modular nucleotide components (1-4bp in length) which each exert a significant effect on the mutation rate of the G_n_T motif. This work advances our ability to accurately identify the position and quantify the mutagenicity of hotspot motifs predicated on short tracts of nucleotides.

## Introduction

Mutation is the ultimate source of genetic variation that facilitates evolution. However, as most mutations are deleterious, species have evolved various prevention and repair mechanisms to decrease the average genomic mutation rate to very low frequencies. Bacterial species have some of the lowest average genomic mutation rates, which are typically around 10^-10^-10^-8^ per nucleotide per generation (Lynch, 2010). However, while the average mutation rate may be very low, this is not true for all mutations. Instead, multiple biases skew the likelihood of observing particular mutations, driving some mutations to happen much more frequently than others. For example, transition bias (Stoltzfus and Norris, 2016) and A:T bias (Hershberg and Petrov, 2010) skew which nucleotide substitutions are more likely to occur, and deletion bias (Kuo and Ochman, 2009) skews DNA toward contraction rather than expansion. As well as biases in the types of mutations that are more common, certain genomic positions also exhibit higher mutation rates than others. For example, genomic macrodomains at intermediate positions exhibit higher mutation rates in multiple bacterial species (Dillon, Sung and Lynch, 2018), as do regions near sites of double-stranded breaks (Shee, Gibson and Rosenberg, 2012). More locally, promoter regions for genes facing away from the replication fork can exhibit an increase in indel rates (Sabari Sankar *et al*., 2016). More locally still, a single nucleotide can have a mutation rate that is hundreds of times higher than a neighbouring nucleotide (Schroeder *et al*., 2016). The primary determinant of this latter phenomenon is the local nucleotide sequence itself.

Local DNA sequence is the major driver of mutation rate heterogeneity throughout genomes, which is the case in both bacterial and in human DNA (Schroeder *et al*., 2016; Liu and Samee, 2023). Nucleotide tracts of just five nucleotides can generate local spikes in mutation rate, commonly referred to as ‘mutational hotspots’. These include recognition sequences for dcm methyltransferase (i.e. CCWGG), which increase C:G→T:A transition rate at the second C position by ∼8-fold (Cherry, 2018). A more potent hotspot motif is a homopolymeric tract or simple sequence repeat (e.g. CCCCC). These are hotspots for indel mutations that cause the tract to either expand or contract by one or more nucleotides. A homopolymer of length five increases indel rates by ∼100-fold, and the hotspot continues to increase in potency the longer the tract (Lee *et al*., 2012). The transition rate of a nucleotide can also vary by ∼400-fold depending on the single nucleotides situated either side of it (Schroeder *et al*., 2016). Identifying which nucleotide sequences generate hotspots is therefore of central importance for predicting mutation rate heterogeneity throughout bacterial genomes. However, as not all nucleotide tracts that generate mutational hotspots are so readily identified, the challenge is to distinguish which nucleotide motifs impact local mutation rates.

In this work, we characterise a novel nucleotide motif that increases T:A→G:C mutation rates in bacteria by up to a factor of ∼1000 when on the leading replicative DNA strand. We show that T:A→G:C substitution mutational hotspots are generated by a ≥7bp motif. The central component of this motif is a homopolymeric tract of G immediately 5’ to a T nucleotide. Similar to indel hotspots, G tract length is strongly positively correlated with increased T:A→G:C mutation rates at the T position when G length is ≥3. However, we also find that whether the hereafter labelled G_n_T tract operates as a potent mutational hotspot is determined by the immediate flanking nucleotides. Rates of mutation are influenced by the 3’ nucleotide following the T and by the 5’ dinucleotide immediately preceding the G tract, making these elements key components of the G_n_T motif. We demonstrate this hotspot motif through experiments with *Pseudomonas fluorescens* complemented by bioinformatic analysis of *Salmonella* species. We combine these data to show that the impact on mutation rate of each G_n_T motif component is similar in our two species. This suggests that the mutation rates for G_n_T motifs determined for this study will be good predictors of hotspot strength in other species of bacteria, particularly *Gammaproteobacteria*.

## Results

### Homopolymeric tracts of guanine with a 3’ thymine are primary components of T:A→G:C mutational hotspots

Recent analysis of numerous species from the NCBI Pathogens database has found that T:A→G:C mutational hotspots in Pseudomonadota (Proteobacteria) and Bacillota (Firmicutes) are generated when a T residue is preceded by tracts of G e.g., GGGGT→GGGGG (Cherry, 2023). The potency of these hotspots increases with the length of the preceding guanine tract, with T:A→G:C rates on average being over 400-fold higher when a T is preceded by 5 or more G’s relative to when it is preceded by 2 or fewer G’s in *Salmonella*. This mutagenic effect is also much more pronounced when the G_n_T motif is on the leading strand of DNA replication (Cherry, 2023).

Prior to this finding, we performed experimental work using an immotile variant of *P. flourescens* SBW25 and observed that, when populations of this bacterium re-evolved motility, this was most often achieved by fixing mutations at a mutational hotspot within the gene *ntrB*. This hotspot, A:T→C:G at position 289, was found to be sensitive to synonymous nucleotide variation, as the introduction of six synonymous changes within 15bp of position 289 removed the hotspot in SBW25 (Horton *et al*., 2021). The local nucleotide context around the hotspot position was therefore demonstrated to be integral to hotspot potency, but the components of the nucleotide motif that drove the hotspot remained unknown.

The gene *ntrB* is natively encoded on the lagging strand, but from the perspective of the leading strand during replication the hotspot nucleotide sequence 5’-3’ is: GGGGT, which mutates to become GGGGG. Therefore, this experimentally determined T:A→G:C mutational hotspot may be an example of the phenomenon observed within the NCBI Pathogens dataset. The first objective of the current study was to determine if guanine tract length was the primary facilitator of the mutational hotspot within *ntrB*, and by doing so provide experimental evidence of the widespread mutational hotspot phenomenon observed throughout many bacterial taxa.

Our approach for measuring changes in mutational hotspot potency was simple: we re-evolved motility in immotile derivatives of *P. fluorescens* SBW25 and measured how often the hotspot mutation was observed. The motility phenotype can be acquired through a single *de novo* mutation from a pool of possible mutations in multiple loci, including the T:A→G:C mutation of interest. We assessed hotspot potency by measuring the proportion of independent replicates that acquired the T:A→G:C mutation as opposed to other viable mutations within the same gene or elsewhere (Fig 1). If the proportion of independent replicates that acquired the T:A→G:C hotspot mutation increased then this showed an increase in mutation rate at the motif, and if it decreased then the reverse was true.

**Fig 1.**
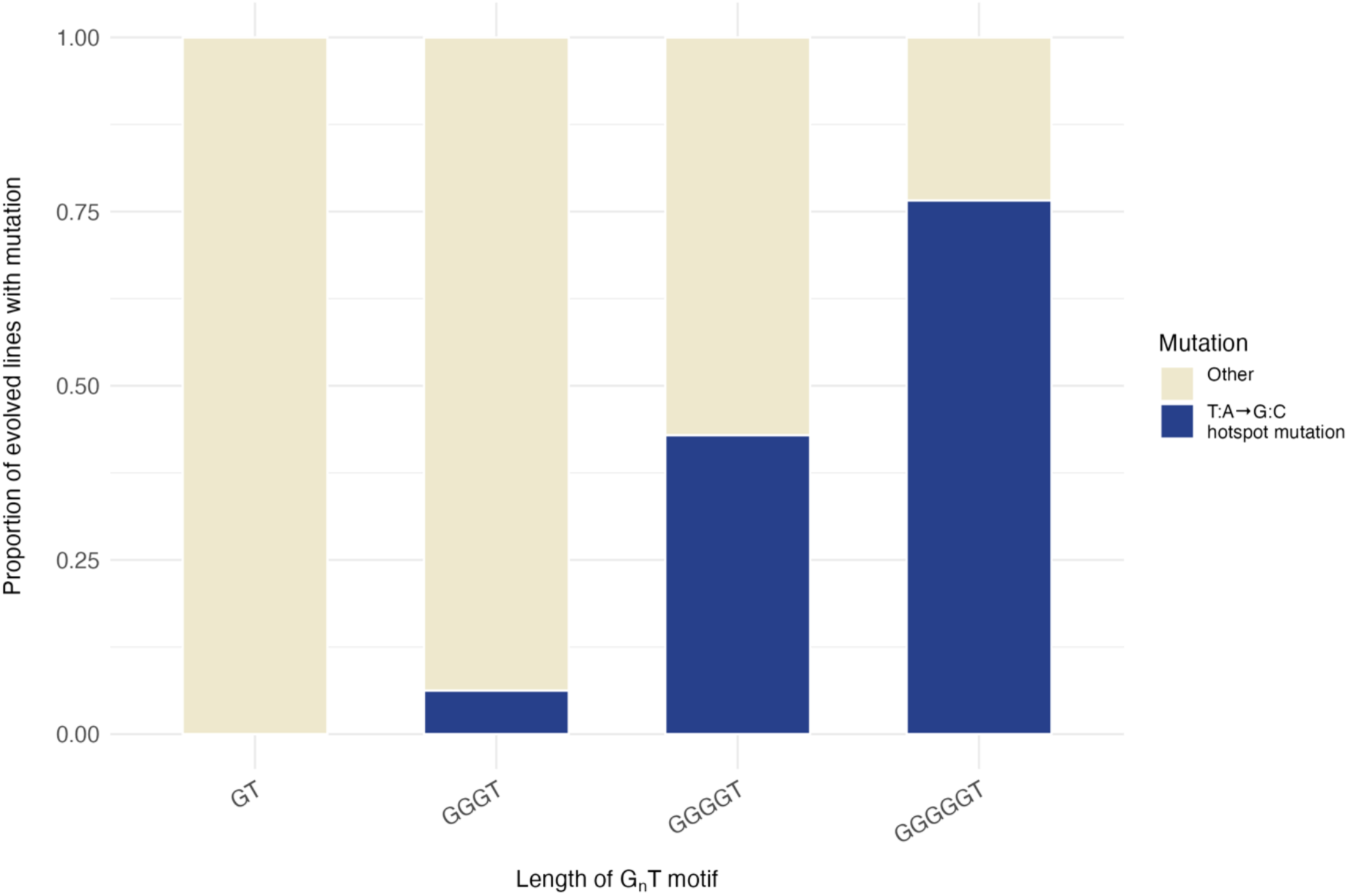
T:A→G:C mutation frequency is positively correlated with length of the preceding G tract. 10 *ntrB* variants with synonymous nucleotide changes neighbouring a T position were used to study the T:A→G:C hotspot motif. Each motif variant contained a homopolymeric tract of guanine 5’ to the T that was either 1, 3, 4, or 5 bp in length. Shown are the aggregated proportion of evolved mutant lines that acquired a T:A→G:C hotspot mutation when the thymine of interest is preceded by 1 G (one strain, *n* = 29), 3 G’s (two strains, *n* = 112), 4 G’s (six strains, *n* = 434), or 5 G’s (one strain, *n* = 47). Total evolved replicates = 622. Overall our experimental work finds that preceding G tract length is a strong determinant of a T:A→G:C mutational hotspot (R^2^ = 0.82).

For all but one strain, the only difference between the hotspot variants evolved for this study were 1-9 synonymous nucleotide variations within the gene *ntrB* that changed the local nucleotide sequence neighbouring the thymine position(s) of interest. Among the synonymous changes introduced were those that changed the length of the preceding G tract. We first pooled the results for all synonymous variant motifs surrounding a mutable T (explained in more detail in the next section) and looked at their hotspot potencies. We observed a clear positive correlation between T:A→G:C hotspot mutation rate and the length of the preceding G tract (Fig 1; R^2^ = 0.82). Our dataset therefore experimentally corroborates the previous analysis in *Salmonella* and additionally demonstrates that the *ntrB* T:A→G:C hotspot is a product of the G_n_T motif.

### T:A→G:C hotspot motifs in experimental populations of P. fluorescens are contingent on preceding G tract length, the flanking 5’ dinucleotide, and flanking 3’ nucleotide

Having verified the importance of homopolymeric tracts of guanine for T:A→G:C mutational hotspots, we next asked whether the motif was affected by other nucleotides in the immediate neighbourhood. We assessed this question in two ways. Firstly, the coding sequence around the *ntrB* 289 hotspot has considerable codon flexibility, which allowed us to probe the impact of the immediate flanking nucleotides. We constructed strains with synonymous substitutions at the second position in the G tract (natively G_4_T), the nucleotide 5’ to the G tract, and the nucleotide 3’ to the mutable T i.e. NGGNGTN (N = any nucleotide). Secondly, we wanted to confirm that any determined sequences exhibited high mutation rates only because of the short tract of nucleotides we augmented, and were not contingent on the wider nucleotide neighbourhood. To do this, we selected an alternative coding A:T position – 683 in *ntrB* – that had been previously shown to also facilitate motility (Horton *et al*., 2021). This position natively has 3 preceding G’s, which we were able to extend synonymously. Additionally, we were able to synonymously mutate the 5’ dinucleotide i.e. SWNGGGTC (S = G/C, W = A/T). In total, we engineered and evolved 10 hotspot motif variants across the two hotspots positions in order to determine each nucleotide component’s impact on hotspot potency (Fig 2).

**Fig 2.**
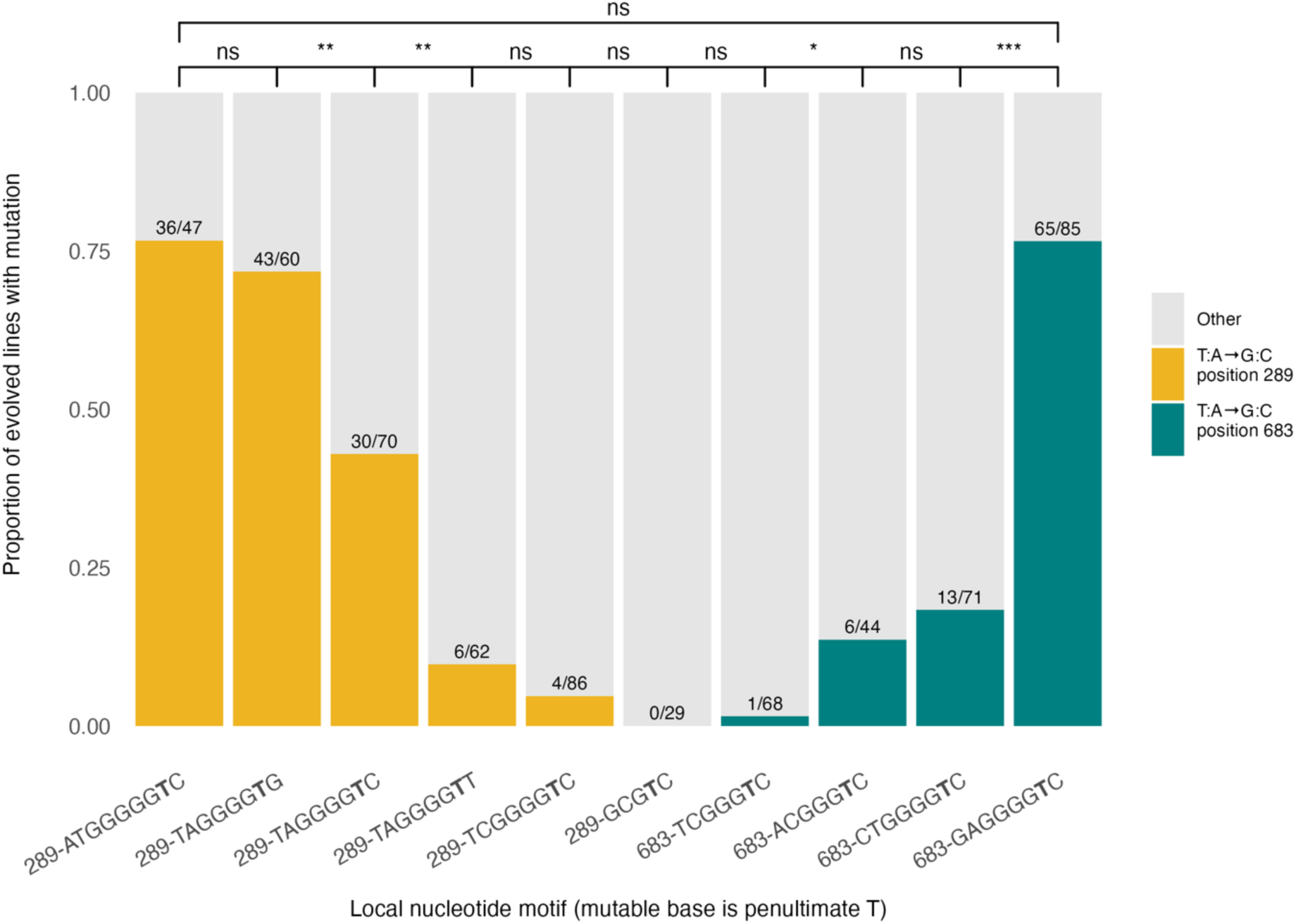
The frequency of T:A→G:C mutation in *P. fluorescens* is strongly influenced by neighbouring G tract length and flanking nucleotide sequence. Mutation frequencies of T:A→G:C are shown for a suite of short synonymous nucleotide tract variants surrounding two T positions within the gene *ntrB*. The sequences on the x-axis, listed 5’-3’ with respect to leading strand replication, show the elements observed to influence mutation rate: the dinucleotide 5’ to the guanine tract, a homopolymeric tract of guanine (1-5bp in length), the hotspot T position that mutates into G (bold), and the nucleotide 3’ to the mutable thymine. T:A→G:C hotspot motifs were augmented around two thymine positions in *ntrB*: coding position 289 (yellow bars) and coding position 683 (teal bars). The y axis shows the proportion of independent populations that acquired the T:A→G:C hotspot mutation, calculated from the observed counts plotted over each bar. Significance values were determined using Fisher’s exact test (* *p* < 0.05; ** *p* < 0.01; *** *p* < 0.001).

Our results reveal that the modular components flanking the core of the hotspot motif – the 5’ dinucleotide and the 3’ nucleotide – exert a significant effect on the mutational hotspot. We evolved three variants of the 289 hotspot that had identical 5’ TA dinucleotides and G tract lengths of 4 (G_4_), with the only difference being the 3’ nucleotide flanking the mutable T (Fig 2; bars 2-4). We observed that hotspot potency significantly decreased as the 3’ nucleotide changed from G to C to T (Fisher’s exact test, p values < 0.0014). Similarly, two motifs around the 289 hotspot that only varied at the 5’ nucleotide position (Fig 2; bars 3 and 5) revealed that a 5’ A is significantly more mutable than a 5’ C (Fisher’s exact test, p << 0.001). Finally, two motifs with a 5’ C, G_3_ tract, and 3’ C surrounding the 683 hotspot significantly differed in hotspot potency when the 5’ dinucleotide partner changed from T to A (Fig 2, bars 7-8; Fisher’s exact text, p < 0.015). The dinucleotide partner was also found to exert a significant effect on motifs of 5’ A, G_4_ tract, and 3’ C around the 289 and 683 hotspots, as G was significantly more mutable than T (Fig 2, bars 3 and 10; p < 0.001).

These results show that the immediate flanking nucleotides are significant components of G_n_T motifs. Additionally, our building of a hotspot at position 683 demonstrates that the motif is not contingent on the wider nucleotide neighbourhood. Outside of the 8bp motif window, the regions around positions 289 and 683 share very little sequence similarity (±6bp contains 2/12 base identity). Yet we were able to increase mutation rate at this position to surpass the native 289 position by augmenting only the established G_n_T motif nucleotides. We observed that hotspot potency was significantly increased around position 683 when changing either the 5’ dinucleotide partner or extending the G tract length to four (Fig 2, bars 7-9; Fisher’s exact text, p < 0.015). The G_4_ variant however remained significantly less mutable than the native 289 hotspot (Fig 2, bars 3 and 9; Fisher’s exact text, p < 0.01) until we changed the 5’ dinucleotide from CT to GA. After this change the 683 hotspot exceeded the native 289 hotspot sequence (Fig 2, bars 3 and 10; Fisher’s exact text, p << 0.001). Therefore, by only augmenting the short nucleotide motif of 8-9bp (depending on G tract length) we are able to build T:A→G:C hotspots within multiple nucleotide neighbourhoods.

### T:A→G:C hotspot motif analysis in natural populations of Salmonella shows that flanking nucleotides determine whether mutations rates are ∼5-fold or >1000-fold above the genomic average

Our experimental assay with *Pseudomonas fluorescens* was able to demonstrate whether a nucleotide substitution within a T:A→G:C hotspot motif exerted a significant effect on hotspot potency. This assay is however limited, primarily by sample size, in its ability to reveal differences between hotspots with low mutation rates and sometimes between those with similar mutation rates. Another limitation is that we were only able to obtain rough estimates of relative mutation rates among motifs. Fortunately, we were able to reinforce our experimental data by re-analysing the *Salmonella* dataset that was generated by (Cherry, 2023) in order to accurately estimate the mutation rate for each unique G_n_T motif. We adapted the original script from (Cherry, 2023) to estimate mutation rates for extended motifs that included the flanking nucleotides. As most of our experimental data focussed on G_4_T motifs we focus on this tract length for our analysis (Figs 3-4), but we also determined T:A→G:C mutation rates for G_3_T and G_5_T motifs (Sup figs 1-2).

**Fig 3.**
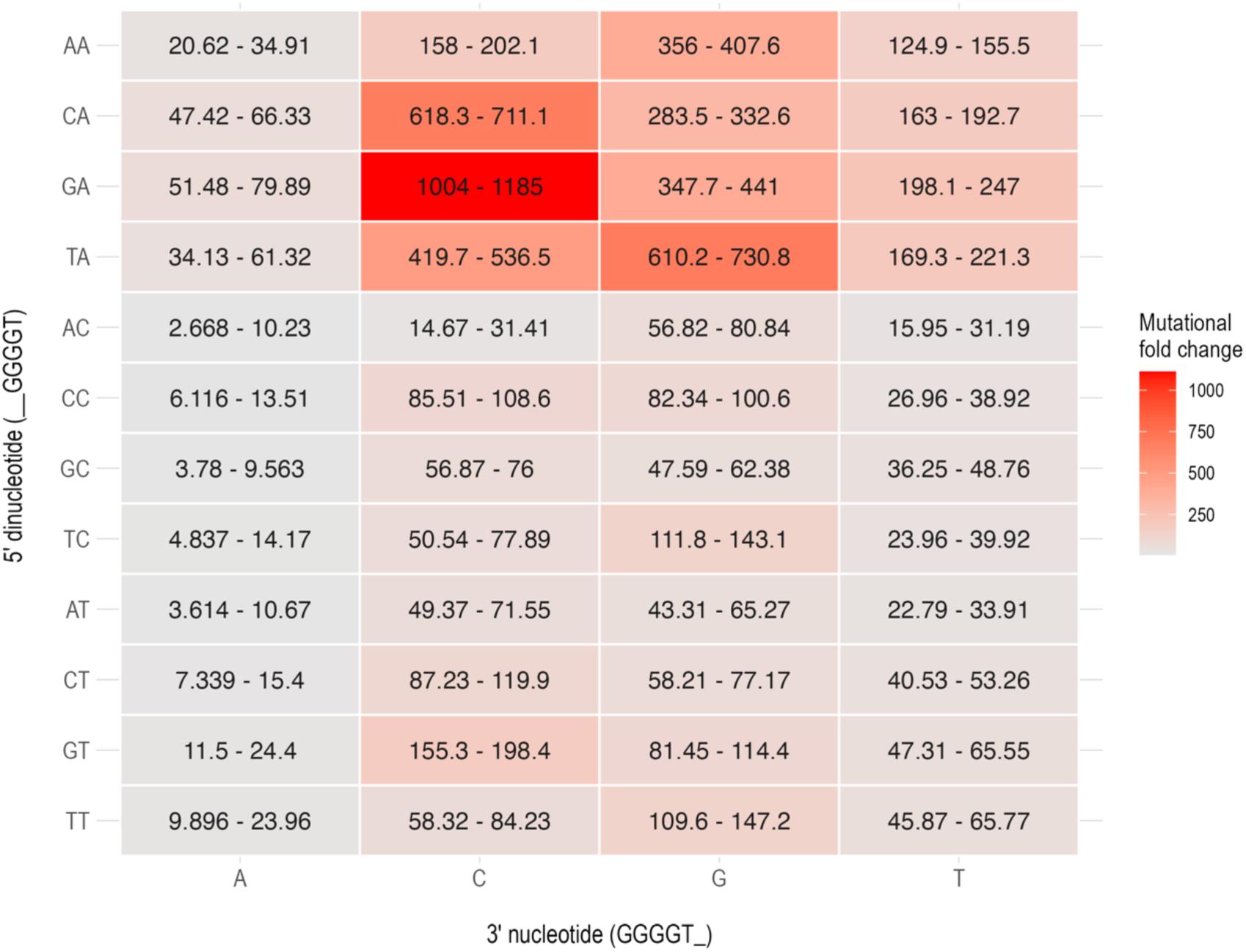
The impact of flanking nucleotides neighbouring G_4_T tracts on T:A→G:C mutation rates in natural *Salmonella* populations. A heatmap showing the T:A→G:C mutation rate for G_4_T motif variants relative to the average T:A→G:C genomic mutation rate with no preceding Gs. Each panel represents a unique combination of nucleotides (5’ dinucleotide on y axis and 3’ nucleotide on x axis) flanking a G_4_T motif. Panels are coloured according to mean mutation rate and annotated with 95% confidence intervals.

**Fig 4.**
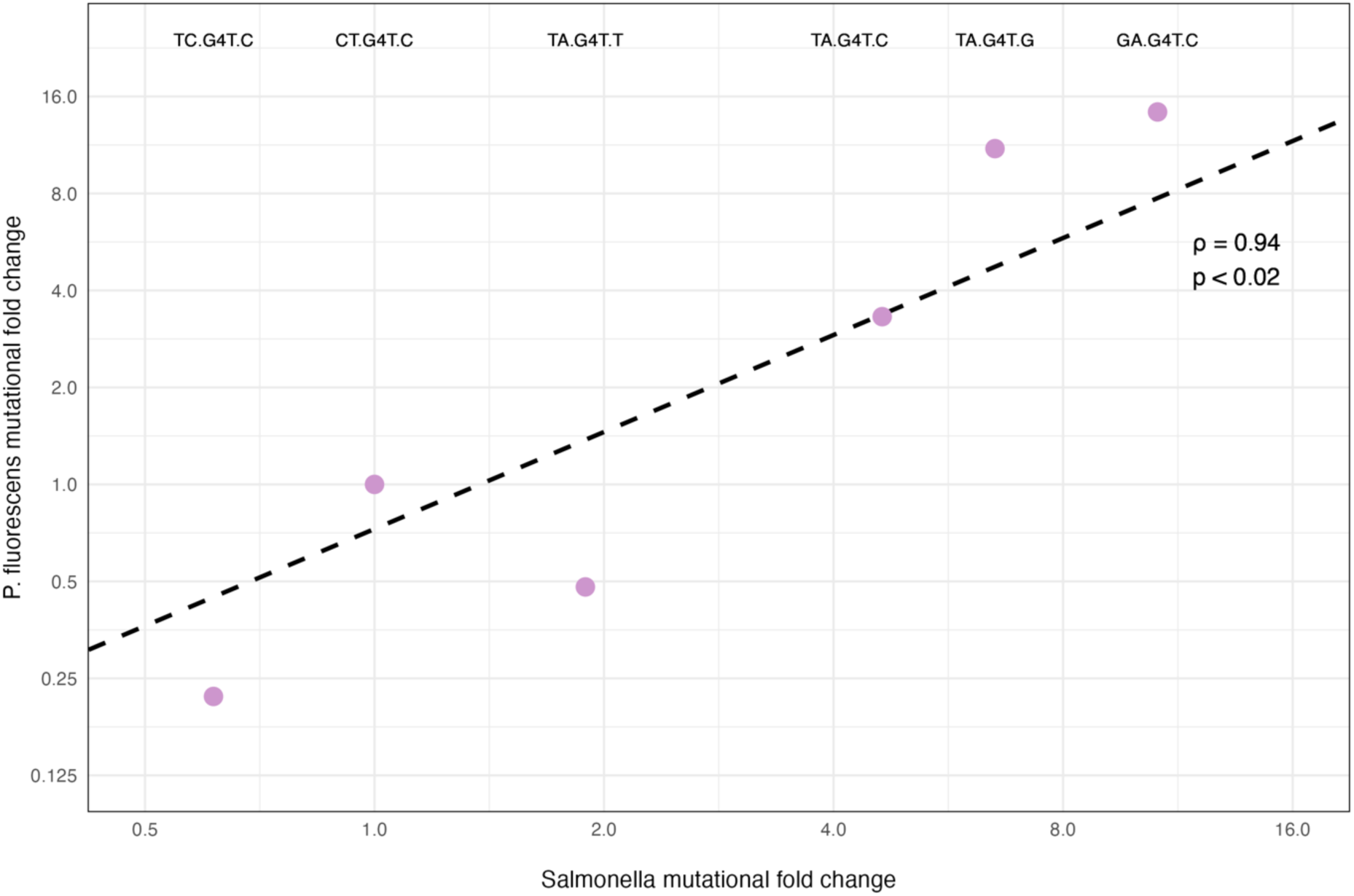
Relative mutation rates of G_4_T motifs in *P. fluorescens* are highly predictive of those in *Salmonella*. An experimental P*. fluorescens* dataset and analysis of natural *Salmonella* populations were made comparable by determining mutational fold-changes of G_4_T motifs (labels) relative to the motif CTG_4_TC (CT.G4T.C). Points show mutational fold changes of *P. fluorescens* motifs (y-axis; determined by Fisher’s exact test odds ratio) and fold changes observed for each motif in *Salmonella* (x-axis; see Methods). The line represents the least squares fit in the log domain. Values for rho and *p* were determined by a Spearman rank-order correlation.

Our analyses found that mutation rates differed by several orders of magnitude depending on the flanking nucleotides. At the lowest end, flanking nucleotides of 5’ AC and 3’ A almost entirely eliminate the G_4_T motif hotspot, with T:A→G:C rates being only ∼5.6-fold higher than genomic T:A→G:C rates when there are no preceding G’s (95% CI = 2.668-10.23; Fig 3). At the high end, flanking nucleotides of 5’ GA and 3’ C lead to T:A→G:C mutation rates ∼1112-fold higher than the genomic average (95% CI = 1004-1185; Fig 3). In general, the mutation rate is highest when the 3’ nucleotide is a C, then G, then T, then A. For the 5’ nucleotide the reverse is true, with highest mutation rates being associated with A, then T, then C.

The 5’ dinucleotide partner’s influence is more variable and demonstrates the context-dependency of motif mutagenicity. For example, 5’ dinucleotide GA typically increases mutation rates most, but TA is the most mutable when the 3’ nucleotide is a G (Fig 3). This finding is corroborated by our experimental data, as the mutation rate of 5’ TA, 3’ C is significantly lower than both 5’ TA, 3’ G and 5’ GA, 3’ C (Fig 2, bars 2, 3 and 10). We additionally observed that the relative impact of the flanking nucleotides on hotspot potency varies depending on G tract length. For motifs with 3’C, the most mutable 5’ dinucleotide is AA for G_3_ (Sup fig 1), GA for G_4_ (Fig 3), and CA or TA for G_5_ (Sup fig 2). Similarly, for motifs with 5’ TA, the most mutable 3’ nucleotides are C then G for G_3_ (Sup fig 1), G then C for G_4_ (Fig 4), and C then T for G_5_ (Sup fig 2).

We next scrutinised the relationship between our datasets by comparing laboratory estimates of relative mutation rates for various motifs in *P. fluorescens* to the estimates obtained for *Salmonella* from analysis of natural isolates (Fig 4). The relative rates correlate well between the two organisms (Spearman’s rank order correlation, rho = 0.94, p < 0.02). This correlation coefficient is expected to be an underestimate due to uncertainty in the rate estimates, especially the large uncertainties observed for our *P. fluorescens* data. Despite this, the two datasets exhibited highly similar mutation rate estimates for the set of motifs we analysed.

The *Salmonella* analyses reinforce the finding that the flanking nucleotides are important determinants of mutation rates at G_n_T motifs, and the data provide accurate quantifications of the flanking nucleotide impact on mutation rates. The analyses also show that the effects of the 5’ dinucleotide, G tract length, and 3’ nucleotide on T:A→G:C mutation rate are not independent. Rather, the effect on mutation rate of changing one component of the motif is contingent on the other components.

## Discussion

Mutation rates across a genome are shaped by multifaceted mechanisms, resulting in significant variation rather than uniformity. Specific sites, known as ‘mutational hotspots,’ exhibit mutation rates much higher than the genomic average and can influence both the direction and predictability of bacterial adaptation (Horton and Taylor, 2023). While it’s known that local nucleotide sequences impact mutation rates (Schroeder *et al*., 2016), the specific features and combinations defining hotspots are not well understood. In this study, we employed experimental and bioinformatic methods to characterise a short nucleotide motif (G_n_T) that can increase T:A→G:C mutation rates by over 1000-fold in bacteria. Through experimental evolution with *P. fluorescens* and complementary bioinformatic analysis of *Salmonella* species, we build on previous work that found that T:A→G:C mutation hotspots are generated by homopolymer G tracts with a 3’ T (Cherry, 2023) to show that G_n_T motifs depend on homopolymeric G tracts in combination with the flanking nucleotides. These findings highlight the modularity of hotspot components, providing insights that could enhance predictive models of mutation rate variability.

### Possible mechanism

The mechanism driving high mutation rates at G_n_T motifs is unclear. However, the central role played by the homopolymeric tract (also referred to as a simple sequence repeat) suggests slipped-strand mispairing as a causal factor. During DNA replication, dissociated parent and nascent DNA strands can re-anneal out of register due to homology with other nucleotides in the repeat region (Levinson and Gutman, 1987). This process typically results in an indel mutation as the out-of-register mispairing causes a nucleotide to loop out of either the nascent or template strand, causing an insertion or deletion of the repetitive nucleotide respectively (Fan and Chu, 2007). In this study we observe a substitution mutation, but this may be a product of slipped-strand mispairing if the T→G mutation is caused by insertion of a G followed by another dissasociation event and re-pairing in the original register, resulting in a G:A mispair. If extension then continues and the mismatch is not repaired, the result will be a substitution of the original T:A hotspot position with its replacement G:C.

It is not clear how the flanking nucleotides contribute to the proposed slipped-strand events, although there are several possibilities. Short nucleotide sequences that generate mutational hotspots during DNA replication must overcome multiple barriers. A sequence must initially exploit error-prone features of the replicating polymerase to generate a mismatch or indel, and subsequently this error must evade proof-reading and mismatch repair to ensure that the mutation becomes immortalised. The flanking nucleotides may contribute to mutagenicity during this process, either by facilitating the slipped-strand mispairing events - by making the DNA sequence more prone to opening or “breathing” (Oron-Karni *et al*., 1997; Jose *et al*., 2009) - or by aiding the resultant error to remain unrecognised by DNA proofreading and repair enzymes.

Proofreading by DNA polymerase discriminates mismatches by checking for correct Watson-Crick geometry (Bębenek and Ziuzia-Graczyk, 2018), likewise mismatch repair recognises errors based on DNA secondary structure (Pavlova *et al*., 2020). Previous works have found direct and indirect evidence that immediate flanking nucleotides can help stabilise mismatches (Allawi and SantaLucia, 1998; Long *et al*., 2018), and that the structure of the DNA helix can be impacted several nucleotides away from a mismatch (Rossetti *et al*., 2015). We estimated melting temperatures for all 48 G_4_T motif sequences assuming a G:A mismatch and when we controlled for the 5’ nucleotide we found a significant positive correlation between DNA stability and mutation rate for the 5’ dinucleotide partner and 3’ nucleotide. Conversely, when we controlled for the 3’ nucleotide we observed a mostly non-significant but negative correlation between stability and mutation rate for the 5’ nucleotide and 5’ dinucleotide partner (Sup fig 3). This relationship can be explained by the preference for C:G/G:C at the 3’ position and against C:G/G:C at the 5’ position. Therefore the flanking nucleotides may impact the stability in different ways, e.g. the low stability of the 5’ dinucleotide may facilitate DNA dissasociation and breathing, while the 3’ nucleotide may stabilise the DNA double helix following a return to an in-register mismatch, helping the error to evade repair. If DNA stability is an important factor in hotspot potency, we can speculate further and note that the purine-purine G•A pair exhibits unique undertwisting of the DNA helix relative to other mismatches (Rossetti *et al*., 2015), which may help explain why the homopolymer substitution phenomenon appears to be unique to G_n_T motifs (Cherry, 2023). Finally, while documentations of sequential indel and mismatch events resulting from strand-slippage are rare, at least one study has previously reported on the phenomenon and implicated DNA breathing in facilitating the effect (Oron-Karni *et al*., 1997).

### Quantifying and controlling mutagenesis

A key advantage to identifying the modular nucleotide components of hotspot motifs and quantifying their impact on hotspot strength is that it allows us to accurately predict mutation rates. It is reassuring that the relative mutation rates of G_n_T motifs determined from *Salmonella* are applicable to *P. fluorescens*, a distantly related species in the *Gammaproteobacteria* class (Williams et al., 2010). This suggests that predicting mutation rates at G_n_T hotspot motifs using the *Salmonella* data will be reasonably accurate across broad genomic backgrounds. However, the potencies of hotspot motifs are additionally influenced by other genomic features such as genomic position and DNA structure (Shepherd, Horton and Taylor, 2022; Liu and Samee, 2023). Thus, it is less certain whether identical motifs within the same genome mutate at the same rates.

The previous *Salmonella* analysis examined rates of T:A→G:C accumulation at G_n_T motifs as a function of chromosome position and found a roughly uniform distribution (Cherry,2023). This suggests that there is no strong systematic dependence of hotspot potency on genome position. However, our previous work found that the *ntrB* G_n_T motif is more potent when situated in its native position as opposed to the Tn7 insertion site, which is situated on the other replichore and is closer to the replication origin (Shepherd et al., 2022). Additionally, recent work has shown that DNA replication error hotspots are affected by multiple genomic properties that fluctuate throughout the genome (Hasenauer *et al*., 2024). Therefore, there may be instances where hotspot potency changes by perhaps an order of magnitude (using the data from Shepherd et al., 2022, as a rough guide) depending on genomic context. However, this phenomenon does not appear to be a simple effect of distance from the origin of replication.

The external genomic parameters influencing the exact mutagenicity of G_n_T motifs are yet to be determined, but our combinatorial data show that the local eight-base pair nucleotide motif is the major determinant of T:A→G:C hotspot mutation rate throughout, and across, bacterial genomes. Therefore, finding these motifs in sequence data can identify the location of potent T:A→G:C hotspots. For example, there are 2078 naturally occuring G_3-5_T motifs on the leading replicative strand of *P. fluorescens* SBW25 with forecast mutation rates >100-fold above the genomic average. Of these, ≥1206 genes (∼20% of reading frames) posess at least one hotspot motif that would result in a non-synonymous T:A→G:C substitution. As well as occurring naturally in many coding sequences, G_n_T motifs are so short that they can also be readily engineered into circuits to help control the evolvability of a system. We investigated this with an experiment where we added two hotspot motifs into the *ntrB* gene, and predictably observed that evolution happened significantly faster and became more predictable at the level of locus (Sup fig 4). There is a growing interest in our ability to both forecast and exert control over the evolutionary process (Lind *et al*., 2019; Castle, Grierson and Gorochowski, 2021). G_n_T motifs, as agents that heavily bias the process of mutation, have the potential to contribute to both of these goals.

## Methods

### Strains and culture conditions

Experiments were performed using populations derived from *Pseudomonas fluorescens* SBW25 strain AR2 (SBW25ΔfleQ IS-ΩKm-hah: PFLU2552) which lacks functionality of FleQ and ViscB, rendering populations immotile (Alsohim *et al*., 2014). The strains constructed from this genomic background contain synonymous nucleotide substitutions in the gene *ntrB*, which has previously been found to resurrect motility under directional selection following a one-step *de novo* mutation in the coding sequence. *P. fluorescens* populations were cryogenically stored at -70°C and grown at 27°C; overnight cultures were agitated at 180rpm. *E. coli* strains used for cloning (DH5α), conjugal transfer (SP50 pRK2073), transposon-mediated genome incorporation (S17 pTNS2), and allelic exchange (ST18) were cryogenically stored at -70°C and grown at 37°C; overnight cultures were agitated at 200rpm.

Antibiotics used in this study were prepared at the following working concentrations: 50 μg/ml kanamycin sulfate; 100 μg/ml ampicillin sodium salt; 30 μg/ml gentamicin sulfate; 10 μg/ml tetracycline hydrochloride; 250 μg/ml streptomycin sulfate salt. All antibiotics were purchased from Sigma-Aldrich.

### Synonymous manipulations of ntrB

To generate synonymous motif variants, we amplified the *ntrBC* operon from *P. fluorescens* SBW25 derivatives (NCBI Assembly: ASM922v1, GenBank sequence: AM181176.4) using outside primers ntrBC-HindIII-F and ntrBC-SacI-R (Sup Table 1). In the first PCR, we generated two fragments using the outside primers in combination with complementary oligonucleotides that had approximately 40bp homology to the nucleotide tract flanking *ntrB* position 289 or 683, depending on the desired construct (Sup Table 1). In the second PCR, we annealed these two fragments with complementary overhangs to generate a complete *ntrBC* operon with synonymous variations at positions neighbouring 289 or 683. For *ntrB* 289 variants we used *P. fluorescens* SBW25 strain AR2 as template DNA; for *ntrB* 683 variants we used *P. fluorescens* SBW25 strain AR2-sm as template DNA to remove the native hotspot at *ntrB* 289. All changes made to the nucleotide sequence left the encoded protein sequence unchanged, both with the original T and with the G resulting from the mutations of interest. Most were single-base changes at third positions of codons. The exception was a change of AGC, which encodes serine, to TCC, which also encodes serine.

### Genomic integration of variant sequences through mini-Tn7 mediated transposition

All *ntrBC* operon variants, aside from 289-TAGGCGTC, were integrated into a non-motile *P. fluorescens* SBW25 derivative AR2 Δ*ntrBC* genomic background using mini-Tn7 mediated transposition (Choi and Schweizer, 2006). As the native operon has been deleted in the host genomic background this incorporation operates as a functional translocation. Generated *ntrBC* inserts and the plasmid pJM220 (pUC18T-miniTn7T-gm-rhaSR-PrhaBAD; a gift from Joanna Goldberg (http://n2t.net/addgene:110559; RRID:Addgene_110559); see Meisner and Goldberg, 2016) were digested with the restriction enzymes HindIII-HF and SacI-HF (NEB) for 1h at 37°C. This was followed by a 30-minute digestion at 37°C where RSAP (NEB) was added to the pJM220 digestion mix. The vector and insert were gel-extracted and purified with a Monarch→ DNA gel extraction kit (NEB) and the fragments were annealed using T4 DNA ligase (NEB) ligation at 16°C for 12h, followed by 4°C hold for ∼4h. The ligated plasmid was transformed into cloning *E. coli* strain DH5α and plated onto LB agar containing ampicillin and gentamicin. Conjugal transfer was performed by preparing 10ml overnight LB broth cultures of the recipient *P. fluorescens* strain (supplemented with kanamycin), DH5α pJM220-*ntrBC* (supplemented with ampicillin and gentamicin), and the helper plasmids SP50 pRK2073 (supplemented with streptomycin) and S17 pTNS2 (supplemented with ampicillin). 700μl of the *P. fluorescens* culture was combined with 100μl of each *E. coli* strain to produce 1ml final volume. The mixed population was pelleted through centrifugation at 8000 rcf for 3 minutes, the supernatant removed, and re-suspended in 1ml fresh LB broth. This process was repeated two additional times to remove residual antibiotic in the culture, before a final re-suspension in 50μl LB broth. This mixed culture was added in a puddle to an LB agar plate containing no antibiotic and left for ∼20-30 minutes in a laminar flow hood to dry. The puddle was incubated at 27°C for a minimum of 5h before an inoculating loop was used to transfer a streak from the centre of the puddle to fresh LB agar supplemented with kanamycin and gentamicin. This plate was incubated at 27°C for 1-2 days until distinct colonies could be observed. Cryogenic clonal stocks were produced by growing one colony overnight at 27°C in the presence of kanamycin and gentamicin and storing the resultant culture in 20% glycerol final volume at -70°C.

### Scarless synonymous augmentation of ntrB through two-step allelic exchange

Synonymous variant 289-TAGGCGTC, which is identical to progenitor strain AR2 except for a C→G at nucleotide 291 on the coding strand of the gene *ntrB*, was created through two-step allelic exchange. This protocol closely followed that outlined by (Hmelo *et al*., 2015), barring amendments outlined in (Horton *et al*., 2021), which were made to optimise the protocol for *Pseudomonas fluorescens*. Primers utilised for this work are outlined in Sup Table 1. In brief, an augmented *ntrB* sequence was produced through two PCR steps as outlined above using AR2 as template data, with the amendment that nested primers ntrB_np_F and ntrB_np_R were used to combine the two fragments and add restriction enzyme recognition sequences in the second PCR. The produced insert, and the vector pTS1, were digested with restriction enzymes BamHI-HF and Hind-III-HF (NEB), gel extracted, and ligated as described above. The ligated plasmid was then transformed into *E. coli* strain ST18 and plated onto LB agar supplemented with 50 μg/ml 5- aminolevulinic acid. The transformed ST18 strain was conjugated with *P. fluorescens* SBW25 derivative AR2 by mixing 750 μl ST18 with 250 μl AR2 and pelleting and puddle mating as outlined above. The conjugative puddle was left to incubate overnight on LB agar supplemented with 50 μg/ml 5-aminolevulinic acid, and a streak was transferred onto LB agar supplemented with tetracycline to select for merodiploid AR2 strains. Two-step recombinants were then selected for by plating on NSLB agar containing +15% w/v sucrose, which selects against merodiploids. Surviving colonies were either *ntrB* C291G mutants or revertants to the AR2 progenitor genotype. Successful mutants were verified by Sanger sequencing of the *ntrB* locus using ntrB_1119_F and ntrB_1119F_R. Cryogenic stocks were then prepared as detailed above.

### Experimental evolution to assess hotspot potency

We used soft agar motility assays to select for mutations that restore motility, including the T:A→G:C mutations at positions 289 and 683 in *ntrB* (from Taylor *et al*., 2015). To prepare soft agar, we poured 30 ml of molten LB broth (Sigma-Aldrich) containing 0.25% agar (Fisher Scientific) into standard 88 mm diameter petri dishes and left these overnight to solidify. The following morning, the petri dishes were placed in a laminar flow hood with the lids removed for 30 minutes to dry. A sterile cocktail stick was then used to transfer a single colony to the soft agar. The stick was held perpendicular to the agar and the colony inserted ∼2 mm. Plates were inoculated in the centre of the dish aside from strains 289-TCGGGGTC, 289-TAGGCGTC, and 683- TCGGGTC, which were inoculated with four equally separated colonies. The plates were incubated at 27°C and monitored daily for emergence of motility zones for ≤12 days. The first motile zone to appear per plate was passaged onto fresh agar containing kanamycin and gentamicin by inserting an inoculating loop into the leading edge of the motile zone and streaking to produce single colony forming units. The motility plate was then discarded to ensure that all emergent motile populations evolved independently.

Passaged plates were incubated for 48 h at 27°C and a single colony was used as template DNA for a colony PCR with primers ntrB_1119_F and ntrB_1119_R (Sup Table 1). The generated *ntrB* amplicons were purified using a Monarch→ PCR DNA clean-up kit (NEB) and Sanger sequencing was performed by Source Bioscience using ntrB_1119_F as a primer. *ntrB* reads provided by Source Bioscience were manually trimmed to remove ∼50bp from each terminus to remove substitution and 1bp indel calls of low confidence. Mutations of high confidence were called by aligning the *ntrB* sequence against the SBW25 reference genomic sequence (NCBI Assembly: ASM922v1, GenBank sequence: AM181176.4) using BLAST.

### Analysis of hotspot motifs in Salmonella species

Estimation of relative mutation rates in natural populations of *Salmonella* was based on the data from Cherry (2023). The number of T→G mutations inferred to have occurred in a sequence context of interest was divided by the number of such sites per genome (a weighted average among many *Salmonella* genomes, as detailed in Cherry (2023)). Confidence intervals were calculated assuming a Poisson sampling distribution for the number of mutations.

### Statistical analysis

Statistical analyses were performed using R version 4.4.1 (2024-06-14). For experimental data, changes in hotspot potency were determined using Fisher’s exact test (fisher.test, from the stats package exact2x2, v4.4.0) with *p* < 0.05 treated as significant. Fold-changes in potency were calculated using Fisher’s exact test’s odds ratio alongside the produced 95% confidence intervals. Figures were generated using R’s *ggplot* package v3.5.1. DNA stability with a G:A mismatch at the mutable T positions was estimated using the R package *rmelting* v1.20.0 (Aravind and Krishna, 2024) with the following parameters: nucleic acid concentration = 2 x 10^-6^, Na concentration = 1.

## References

Allawi, H.T. and SantaLucia, J. (1998) ‘Nearest-neighbor thermodynamics of internal A.C mismatches in DNA: sequence dependence and pH effects’, Biochemistry, 37(26), pp. 9435– 9444. Available at: 10.1021/bi9803729.

Alsohim, A.S. et al. (2014) ‘The biosurfactant viscosin produced by Pseudomonas fluorescens SBW25 aids spreading motility and plant growth promotion’, environmental microbiology, 16(7), pp. 2267–2281.

Aravind, J. and Krishna, G. (2024) ‘rmelting: R Interface to MELTING 5’. Available at: https://aravind-j.github.io/rmelting/.

Bębenek, A. and Ziuzia-Graczyk, I. (2018) ‘Fidelity of DNA replication-a matter of proofreading’, Current Genetics, 64(5), pp. 985–996. Available at: 10.1007/s00294-018-0820-1.

Castle, S.D., Grierson, C.S. and Gorochowski, T.E. (2021) ‘Towards an engineering theory of evolution’, Nature Communications, 12(1), p. 3326. Available at: 10.1038/s41467-021-23573-3.

Cherry, J.L. (2018) ‘Methylation-Induced Hypermutation in Natural Populations of Bacteria’, Journal of Bacteriology, 200(24), pp. e00371–18.

Cherry, J.L. (2023) ‘T Residues Preceded by Runs of G Are Hotspots of T→G Mutation in Bacteria’, Genome Biology and Evolution, 15(6), p. evad087. Available at: 10.1093/gbe/evad087.

Choi, K.H. and Schweizer, H.P. (2006) ‘mini-Tn7 insertion in bacteria with single attTn7 sites: Example Pseudomonas aeruginosa’, Nature Protocols, 1(1), pp. 153–161. Available at: 10.1038/nprot.2006.24.

Dillon, M.M., Sung, W. and Lynch, M. (2018) ‘Periodic Variation of Mutation Rates in Bacterial Genomes’, mBio, 9(4), pp. 1–15.

Fan, H. and Chu, J.-Y. (2007) ‘A Brief Review of Short Tandem Repeat Mutation’, Genomics, Proteomics & Bioinformatics, 5(1), pp. 7–14. Available at: 10.1016/S1672-0229(07)60009-6.

Hasenauer, F.C. et al. (2024) ‘Genome-wide mapping of spontaneous DNA replication error- hotspots using mismatch repair proteins in rapidly proliferating Escherichia coli’, Nucleic Acids Research [Preprint]. Available at: 10.1093/nar/gkae1196.

Hershberg, R. and Petrov, D.A. (2010) ‘Evidence That Mutation Is Universally Biased towards AT in Bacteria’, PLOS Genetics, 6(9), p. e1001115. Available at: 10.1371/journal.pgen.1001115.

Hmelo, L.R. et al. (2015) ‘Precision-engineering the Pseudomonas aeruginosa genome with two- step allelic exchange’, Nature Protocols, 10(11), pp. 1820–1841. Available at: 10.1038/nprot.2015.115.

Horton, J.S. et al. (2021) ‘A mutational hotspot that determines highly repeatable evolution can be built and broken by silent genetic changes’, Nature Communications, 12(6092), pp. 1–10. Available at: 10.1038/s41467-021-26286-9.

Horton, J.S. and Taylor, T.B. (2023) ‘Mutation bias and adaptation in bacteria’, Microbiology, 169(11), p. 001404. Available at: 10.1099/mic.0.001404.

Jose, D. et al. (2009) ‘Spectroscopic studies of position-specific DNA “breathing” fluctuations at replication forks and primer-template junctions’, Proceedings of the National Academy of Sciences, 106(11), pp. 4231–4236. Available at: 10.1073/pnas.0900803106.

Kuo, C.-H. and Ochman, H. (2009) ‘Deletional Bias across the Three Domains of Life’, Genome Biology and Evolution, 1, p. 145. Available at: 10.1093/gbe/evp016.

Lee, H. et al. (2012) ‘Rate and molecular spectrum of spontaneous mutations in the bacterium Escherichia coli as determined by whole-genome sequencing’, 109(41). Available at: https://doi.org/10.1073/pnas.1210309109/-/DCSupplemental.www.pnas.org/cgi/doi/10.1073/pnas.1210309109.

Levinson, G. and Gutman, G.A. (1987) ‘Slipped-strand mispairing: a major mechanism for DNA sequence evolution’, Molecular Biology and Evolution, 4(3), pp. 203–221. Available at: 10.1093/oxfordjournals.molbev.a040442.

Lind, P.A. et al. (2019) ‘Predicting mutational routes to new adaptive phenotypes’, eLife, 8, p. e38822. Available at: 10.7554/eLife.38822.

Liu, Z. and Samee, M.A.H. (2023) ‘Structural underpinnings of mutation rate variations in the human genome’, Nucleic Acids Research, 51(14), pp. 7184–7197. Available at: 10.1093/nar/gkad551.

Long, H. et al. (2018) ‘Specificity of the DNA Mismatch Repair System (MMR) and Mutagenesis Bias in Bacteria’, Molecular Biology and Evolution, 35(10), pp. 2414–2421. Available at: 10.1093/molbev/msy134.

Lynch, M. (2010) ‘Evolution of the mutation rate’, Trends in genetics : TIG, 26(8), pp. 345–352. Available at: 10.1016/j.tig.2010.05.003.

Meisner, J. and Goldberg, J.B. (2016) ‘The Escherichia coli rhaSR-PrhaBAD inducible promoter system allows tightly controlled gene expression over a wide range in Pseudomonas aeruginosa’, Applied and Environmental Microbiology, 82(22), pp. 6715–6727. Available at: 10.1128/AEM.02041-16.

Oron-Karni, V. et al. (1997) ‘A Novel Mechanism Generating Short Deletion/Insertions Following Slippage is Suggested by a Mutation in the Human α2-Globin Gene’, Human Molecular Genetics, 6(6), pp. 881–885. Available at: 10.1093/hmg/6.6.881.

Pavlova, A.V. et al. (2020) ‘Responses of DNA Mismatch Repair Proteins to a Stable G- Quadruplex Embedded into a DNA Duplex Structure’, International Journal of Molecular Sciences, 21(22), p. 8773. Available at: 10.3390/ijms21228773.

Rossetti, G. et al. (2015) ‘The structural impact of DNA mismatches’, Nucleic Acids Research, 43(8), pp. 4309–4321. Available at: 10.1093/nar/gkv254.

Sabari Sankar, T., et al. (2016) ‘The Nature of Mutations Induced by Replication-Transcription Collisions’, Nature, 535(7610), pp. 171–181. Available at: 10.1016/j.physbeh.2017.03.040.

Schroeder, J.W. et al. (2016) ‘The effect of local sequence context on mutational bias of leading and lagging strand genes’, Current Biology, 26(5), pp. 692–697. Available at: 10.1016/j.cub.2016.01.016.The.

Shee, C., Gibson, J.L. and Rosenberg, S.M. (2012) ‘Two Mechanisms Produce Mutation Hotspots at DNA Breaks in Escherichia coli’, Cell Reports, 2(4), p. 714. Available at: 10.1016/j.celrep.2012.08.033.

Shepherd, M.J., Horton, J.S. and Taylor, T.B. (2022) ‘A near-deterministic mutational hotspot in Pseudomonas fluorescens is constructed by multiple interacting genomic features’, Molecular Biology and Evolution, 39(6).

Stoltzfus, A. and Norris, R.W. (2016) ‘On the Causes of Evolutionary Transition:Transversion Bias’, Molecular Biology and Evolution, 33(3), pp. 595–602. Available at: 10.1093/molbev/msv274.

Taylor, T.B. et al. (2015) ‘Evolutionary resurrection of flagellar motility via rewiring of the nitrogen regulation system’, Science, 347(6225), pp. 1014–1017. Available at: 10.1126/science.1259145.

