## Supplementary information for "G_n_T motifs: short nucleotide tracts of ≥8bp that can increase T:A→G:C mutation rates >1000-fold in bacteria"

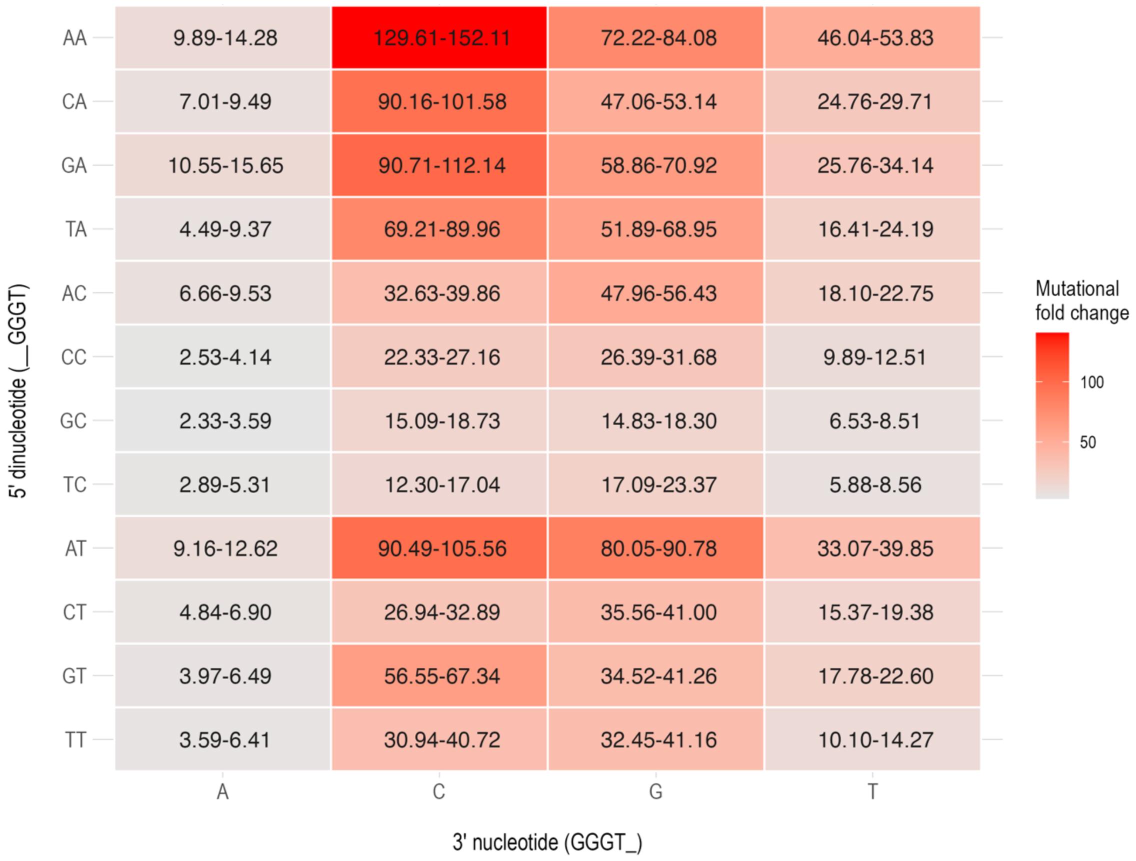


**Sup fig 1. The impact of flanking nucleotides neighbouring $G_3T$ tracts on T:A→G:C mutation rates in natural *Salmonella* populations.** 95 percent confidence intervals are annotated within each cell.

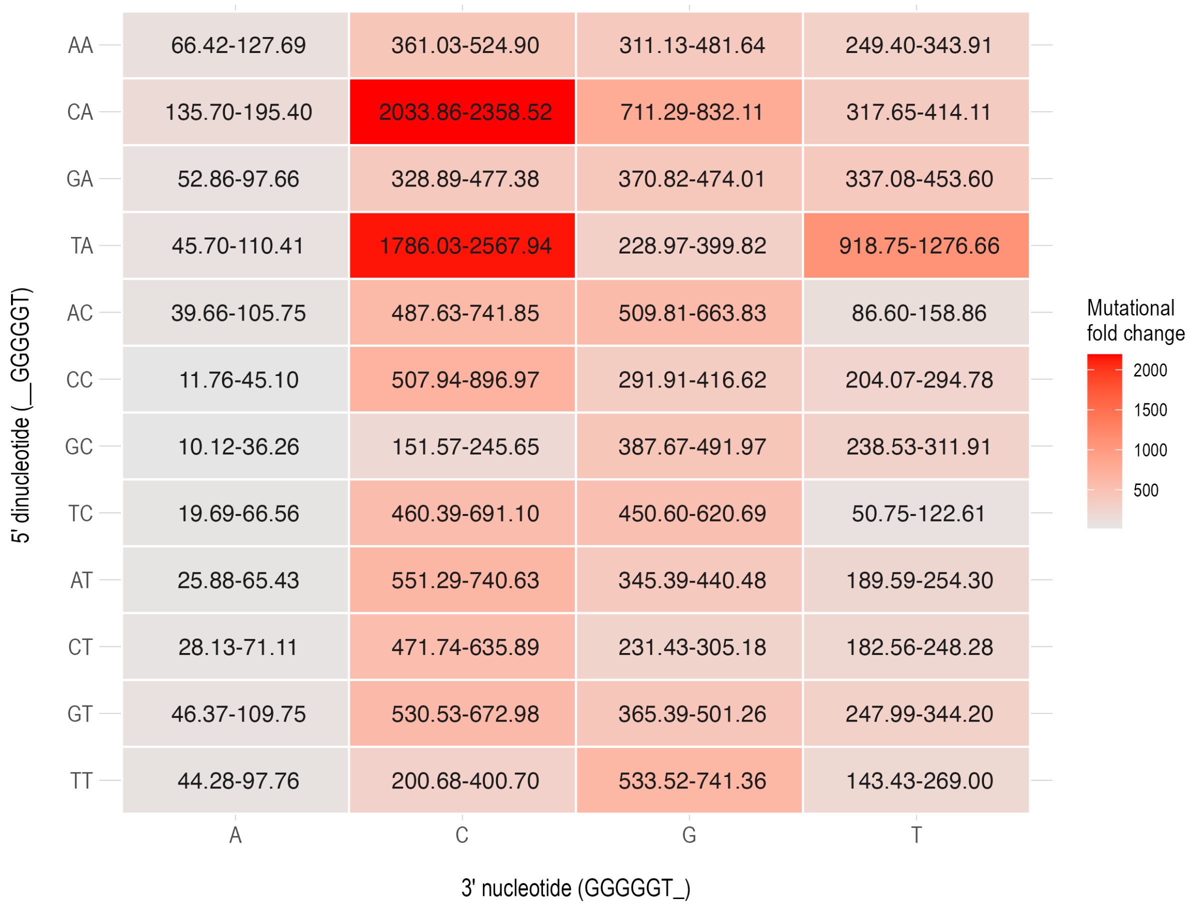


**Sup fig 2. The impact of flanking nucleotides neighbouring $G_5T$ tracts on T:A→G:C mutation rates in natural *Salmonella* populations.** 95 percent confidence intervals are annotated within each cell.

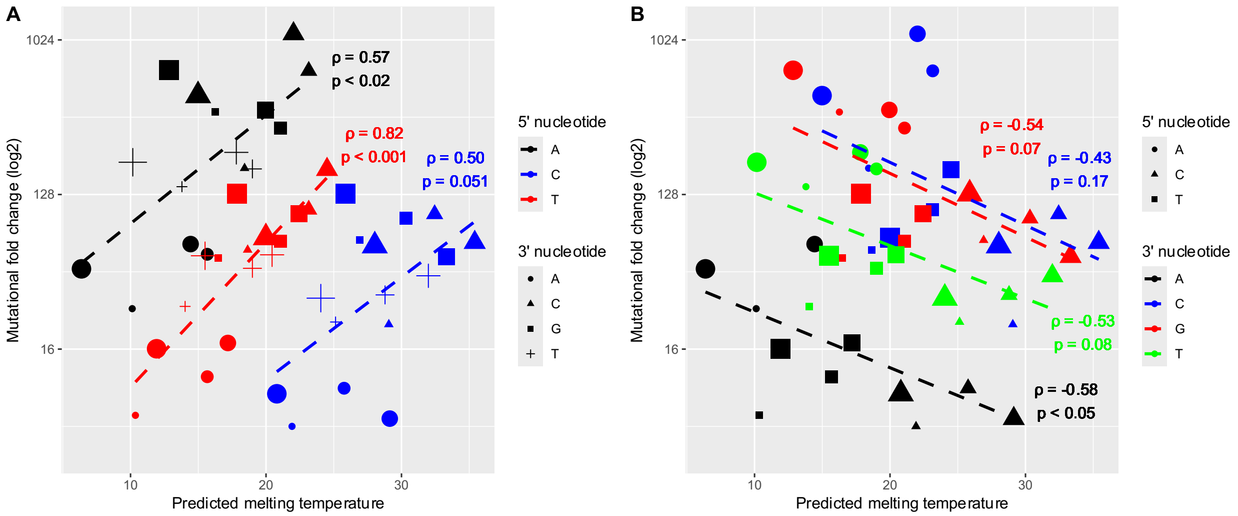


**Sup fig 3. Correlation between melting temperature and T:A→G:C mutation rates of $G_4T$ motifs.** Mutational fold changes (y-axis; values derived from Fig 3) and predicted melting temperatures assuming a G:A mismatch (x-axis; approximated by nearest-neighbour formulations) were plotted for all $G_4T$ motif variants. rho and p values are plotted next to each determined slope when grouped by the (A) 5’ nucleotide, and (B) 3’ nucleotide. 5’ dinucleotide partner nucleotides are denoted by size in ascending order: A, C, G, T.

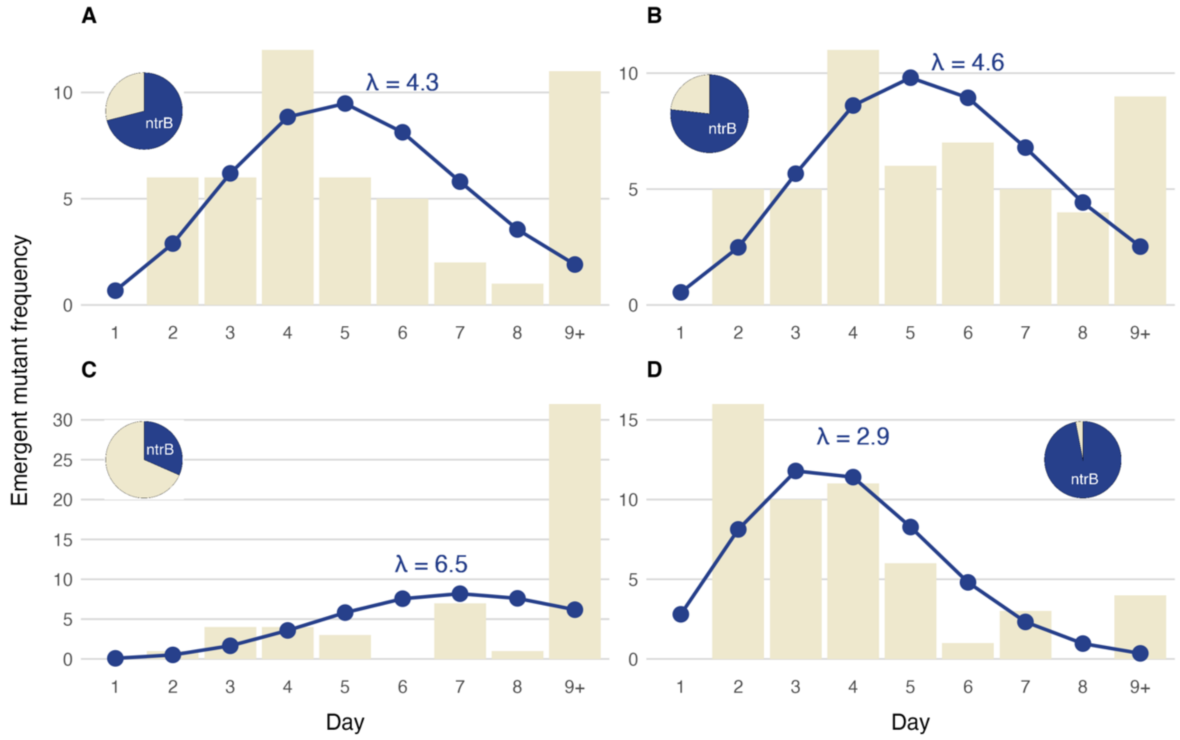


**Sup fig 4. The number and orientation of T:A→G:C hotspot motifs determine rate of evolution and evolutionary predictability.** Emergence of the motility phenotype and the proportion of evolved replicates with *de novo* mutation in *ntrB* were determined for four hotspot motif *ntrB* variants: (A) 289-AT.$G_5$T.C encoded on the leading strand; (B) 683-GA.$G_4$T.C encoded on the leading strand; (C) 289-AT.$G_5$T.C and 683-GA.$G_4$T.C encoded on the lagging strand; (D) 289-AT.$G_5$T.C and 683-GA.$G_4$T.C encoded on the leading strand. Each variant was used to seed a batch of independent replicates (*n* = 49, 52, 52, and 51 respectively). These were placed under selection for motility and monitored for eight days (x-axis). The frequency of the emergent motility phenotype was recorded (y-axis) and replicates that did not evolve within the experimental window were designated to have evolved in 9+ days. Mean emergence times were determined by plotting a Poisson distribution. Replicates that evolved within eight days were sent for Sanger sequencing of the *ntrB* locus (*n* = 38, 43, 19, and 44). The proportion of replicates that acquired a *de novo* mutation in *ntrB* are shown by the pie charts within each panel.

### Sup table 1. Oligonucleotides used in the study.

| Primer name: | Sequence (5'-3'): | Primer use: |
|---|---|---|
| ntrBC-HindIII-F<br>ntrBC-SacI-R | AATTTAAGCTTCACTGTCCGAACAACACTGATC<br>AATTTGAGCTCCGGTTCATGGTGCATTGAAGC | Outside primers used to generate synonymous ntrBC variant inserts. |
| GTT@96-F<br>GTT@96-R | CTGACGGTCGACTACGCCGTTACCCCTATCCTGAGCAACG<br>CGTTGCTCAGGATAGGGGTAACGGCGTAGTCGACCGTCAG | Inside primer pairs combined with ntrBC-HindIII-F and ntrBC-SacI-R to produce 3' motif variants at ntrB position 289. |
| GTC@96-F<br>GTC@96-R | CTGACGGTCGACTACGCCGTCACCCCTATCCTGAGCAACG<br>CGTTGCTCAGGATAGGGGTGACGGCGTAGTCGACCGTCAG | |
| CCC@98-F<br>CCC@98-R | CTGACGGTCGACTACGCCGTGACCCCCATCCTGAGCAACG<br>CGTTGCTCAGGATGGGGGTCACGGCGTAGTCGACCGTCAG | Inside primer pairs combined with ntrBC-HindIII-F and ntrBC-SacI-R to produce 5' motif variants at ntrB position 289. |
| CCG@98-F<br>CCG@98-R | CTGACGGTCGACTACGCCGTGACCCCGATCCTGAGCAACG<br>CGTTGCTCAGGATCGGGGTCACGGCGTAGTCGACCGTCAG | |
| TCC@230-F<br>TCC@230-R | TCACCTTGGTGCGCGACTACGACCCGTCCATTCCCGACGTATTG<br>CAATACGTCGGGAATGGACGGGTCGTAGTCGCGCACCAAGGTGA | Inside primer pairs combined with ntrBC-HindIII-F and ntrBC-SacI-R to produce a 5' motif variant at ntrB position 683. |
| CCC@229-F<br>CCC@229-R | TCACCTTGGTGCGCGACTACGACCCCAGCATTCCCGACGTATTG<br>CAATACGTCGGGAATGCTGGGGTCGTAGTCGCGCACCAAGGTGA | Inside primer pairs combined with ntrBC-HindIII-F and ntrBC-SacI-R to produce a G tract variant at ntrB position 683. |
| CCCTCC@229-230-F<br>CCCTCC@229-230-R | TCACCTTGGTGCGCGACTACGACCCCTCCATTCCCGACGTATTG<br>CAATACGTCGGGAATGGAGGGGTCGTAGTCGCGCACCAAGGTGA | Inside primer pairs combined with ntrBC-HindIII-F and ntrBC-SacI-R to produce a G tract and 5' variant at ntrB position 683. |
| ntrB_Up_F<br>ACG@97-F | GAGAGCTGACCGTTGAAACC<br>CTGACGGTCGACTACGCCGTGACGCCTATCCTGAGCAACG | Primer pair used to generate 5' segment of augmented ntrB locus for allelic exchange. |
| ACG@97-R<br>ntrB_Dn_R | CGTTGCTCAGGATAGGCGTCACGGCGTAGTCGACCGTCAG<br>ATGGCGCGAAACACTTCCTG | Primer pair used to generate 3' segment of augmented ntrB locus for allelic exchange. |
| ntrB_np_F<br>ntrB_np_R | AATTTGGATCCATGACCATCAGCGATGCACTG<br>AATTTAAGCTTGATCCAGACGGTTTCACTACG | Nested primers used to produce complete augmented ntrB locus for allelic exchange. |
| ntrB_1119_F<br>ntrB_1119_R | GAGGTCCCAATGACCATCAG<br>GACGATCCAGACGGTTTCAC | Primer pair used for amplicon production of ntrB. |
